# Multilevel integrative transcriptome analyses in humans and humanized mice define in vivo human lncRNA metabolic regulators

**DOI:** 10.1101/2020.01.01.884023

**Authors:** Xiangbo Ruan, Ping Li, Yonghe Ma, Cheng-fei Jiang, Yi Chen, Yu Shi, Nikhil Gupta, Fayaz Seifuddin, Mehdi Pirooznia, Hiroshi Suemizu, Yasuyuki Ohnishi, Nao Yoneda, Megumi Nishiwaki, Haiming Cao

## Abstract

A growing number of long non-coding RNAs (lncRNAs) have emerged as vital metabolic regulators in research animals suggesting that lncRNAs could also play an important role in human metabolism. However, most human lncRNAs are non-conserved, vastly limiting our ability to identify human lncRNA metabolic regulators (hLMRs). As the sequence-function relation of lncRNAs has yet to be established, the identification of lncRNA metabolic regulators in animals often relies on their regulations by experimental metabolic conditions. But it is very challenging to apply this strategy to human lncRNAs because well-controlled human data are much limited in scope and often confounded by genetic heterogeneity. In this study, we establish an efficient pipeline to identify putative hLMRs that are metabolically sensitive, disease-relevant, and population applicable. We first progressively processed human transcriptome data to select human liver lncRNAs that exhibit highly dynamic expression in the general population, show differential expression in a metabolic disease population, and response to dietary intervention in a small disease cohort. We then experimentally demonstrated the responsiveness of selected hepatic lncRNAs to defined metabolic milieus in a liver-specific humanized mouse model. Furthermore, by extracting a concise list of protein-coding genes that are persistently correlated with lncRNAs in general and metabolic disease populations, we predicted the specific function for each hLMR. Using gain- and loss-of-function approaches in humanized mice as well as ectopic expression in conventional mice, we were able to validate the regulatory role of one non-conserved hLMR in cholesterol metabolism. Mechanistically, this hLMR binds to an RNA-binding protein, PTBP1, to modulate the transcription of cholesterol synthesis genes. In summary, our study provides a pipeline to overcome the variabilities intrinsic to human data to enable the efficient identification and functional definition of hLMRs. The combination of this bioinformatic framework and humanized murine model will enable broader systematic investigation of the physiological role of disease-relevant human lncRNAs in metabolic homeostasis.

## Introduction

In the past two decades, metabolic diseases have reached epidemic proportions globally^1, 2^, underscoring the urgent need for a better understanding of the metabolism regulation in humans. While significant progress has been made in mapping the makeup and wiring diagram of metabolic pathways in recent years, substantial gaps remain in our understanding of the underlying pathophysiology of major metabolic diseases, such as obesity, NAFLD and type 2 diabetes. As maintenance of metabolic homeostasis requires high-level coordination at cellular, organ and organismal levels^3–5^, such level of complexity often cannot be adequately modeled by cultured cells and need to be studied in in vivo systems, such as mice, which have the advantage of being conducive to genetic manipulation. Although these animal-based experimental studies are essential to capture mechanistic and physiologically relevant understanding of the role of metabolism in health and disease, there are limitations due to species differences and the imperfect disease models. These species-distinct differences are further underscored by our incomplete understanding of the human genome, particularly the enormous landscape of non-coding regions. As 2% of the human genome is sufficient to encode all protein-coding genes, the vast majority of the genome is non-coding and was once considered to be made of gene deserts. It is now well established that most of the “non-coding” regions can be transcribed, giving rise to approximately 60,000 lncRNAs^6^, which would equate to three times the number of protein-coding genes. Growing evidence supports that lncRNAs play a regulatory role in systemic energy metabolism in mice. For examples, we have shown that a liver-enriched lncRNA, lncLSTR, regulates systemic lipid metabolism^7^, and a second lncRNA, lncLGR, regulates glycogen content in mice^8^. Robust mouse lncRNA metabolic regulators (mLMRs), such as Lexis, Mexis, and Blnc1^9–11^, have also been reported by many groups and this list continues to expand. Furthermore, several genome-wide transcriptome analyses in mice have identified hundreds of potential mLMRs in key metabolic organs suggesting that mouse lncRNAs constitute an additional dimension of metabolic regulation^12, 13^. If human lncRNAs exercise a similar function, studying their metabolic function could help systemically uncover novel regulatory mechanisms of human metabolism and expand our understanding of how metabolic disease is initiated and progresses.

Despite their enormous potential, it is currently extremely difficult to define human lncRNA metabolic regulators (hLMRs), in part, due to the multitude of challenges in assigning functions to lncRNAs in general, and in determining the metabolic function of human lncRNAs in particular. Current knowledge and technology limit our ability to identify and characterize lncRNA functions especially relative to the progress in protein-coding genes. Since our current understanding of the sequence-function relationship of lncRNAs is very poor, we cannot use sequence features such as functional domains in protein-coding genes, to place lncRNAs in a biological context^14^. To address this challenge in the context of energy metabolism, considerable efforts have been devoted to identifying mLMRs by analyzing the regulatory information of a lncRNA in response to various conditions to inform its function. For examples, we have developed a pipeline to identify mLMRs based on their regulations by multiple pathophysiologically representative metabolic conditions in mice^12^. Sallam et al. identified two mLMRs of cholesterol metabolism, based on their regulations by liver X receptor (LXR), a well-established transcription factor in cholesterol homeostasis^10, 11^. The valuable information yielded by these extensive studies could in theory help identify hLMRs if lncRNAs were as conserved as mRNAs. Surprisingly and intriguingly, however, over 80% of human lncRNAs are not conserved^15, 16^, and most human lncRNAs cannot be found in mice and vice versa. Thus, most human lncRNAs belong to a unique class of molecules whose regulatory information has to be directly derived from human. Although clinical RNA-seq data of human metabolic tissues are emerging and being accumulated, their numbers and the metabolic conditions under which they are collected are limited. More importantly, unlike data generated in inbred mouse strains under well-controlled experimental conditions, the gene expression levels in humans are significantly affected by genetic heterogeneity^17^ and environmental factors. These challenges are further exacerbated by the lack of sequence-based functional inference, which has been routinely used to corroborate the disease relevance of protein-coding genes. With so many complicating factors involved, it is evidently not a trivial task to retrieve hLMR signals from human data that truly reflect metabolic responses, or are metabolically sensitive, without losing their significance to the general population, and an approach that is specifically suitable for human lncRNAs is needed. Furthermore, for protein-coding genes, the definitive functional validation is routinely carried out in research animals, particularly in mice, by creating gain-or loss-of-function models. As most human lncRNAs are non-conserved, their physiological function cannot be directly studied in conventional mice, and an in vivo model is needed to experimentally characterize putative hLMRs in a physiologically relevant setting.

In order to leverage the accumulating clinical studies to understand the pathophysiological importance of lncRNAs in human metabolism, we have established an effective strategy to retrieve a list of broadly representative and metabolically sensitive human lncRNAs from human transcriptome data. We further refined our selection based on the regulation of these human lncRNAs by defined metabolic conditions in a humanized mouse model, and most importantly, experimentally defined the in vivo role of a non-conserved hLMR in cholesterol metabolism in the humanized mouse model.

## Results

### Identification of human lncRNA metabolic regulators

To take advantage of the currently available human data while overcoming their limitations, in this study, we combined a variety of human studies as well as a humanized mouse model to establish a practical platform to identify population applicable, metabolically sensitive and disease-relevant human lncRNA metabolic regulators (hLMRs) (Figure 1). In order to maintain the broad representation of selected lncRNAs in the general population, we started our analysis with human liver RNA-seq data from the Genotype-Tissue Expression (GTEx) project^18, 19^. While GTEx RNA-seq data harbor valuable information for distinct tissue-relevant genes, they do not have deep clinical phenotyping for identifying differentially expressed genes linked to metabolic disease or therapy. Interestingly, information of gene expression variability or dynamics in a specific cell type or tissue has been utilized to infer potential roles in pathophysiology and diseases^20–22^. We postulate that lncRNAs with high expression variability in key metabolic tissues such as liver in the general population from GTEx could be potentially metabolically sensitive and disease-relevant. Therefore, we determined the gene expression variability in the livers of individuals in GTEx. To extend the coverage for human lncRNAs, we have recently established a most updated and comprehensive lncRNA database, lncRNA Knowledgebase (lncRNAKB) (BioRxiv: 669994), which we used to map human lncRNAs throughout this study (see Methods). In total, the coefficients of variation for 16906 genes including 2665 lncRNAs expressed in the liver samples (Figure 2A) were calculated and ranked in four quartiles (Table S1). We then assigned the protein-coding genes in the top and bottom quartile separately to the category of complex diseases using DAVID gene functional annotation tool. As shown in Figure 2B, there is a significant enrichment of multiple disease categories in the top quartile of protein-coding genes from liver, while very few were found in the bottom quartile. This result suggests that hepatic genes with high dynamic expression in the general human population might be conditionally responsive and susceptible to a variety of complex disease conditions particularly cardiometabolic diseases (Figure 2B). Thus the 943 lncRNAs included in the top quartile (Table S1) could be metabolically sensitive and potentially function as human lncRNA regulators in the development of cardiometabolic diseases (Figure 1, Identification step 1).

**Figure 1.**
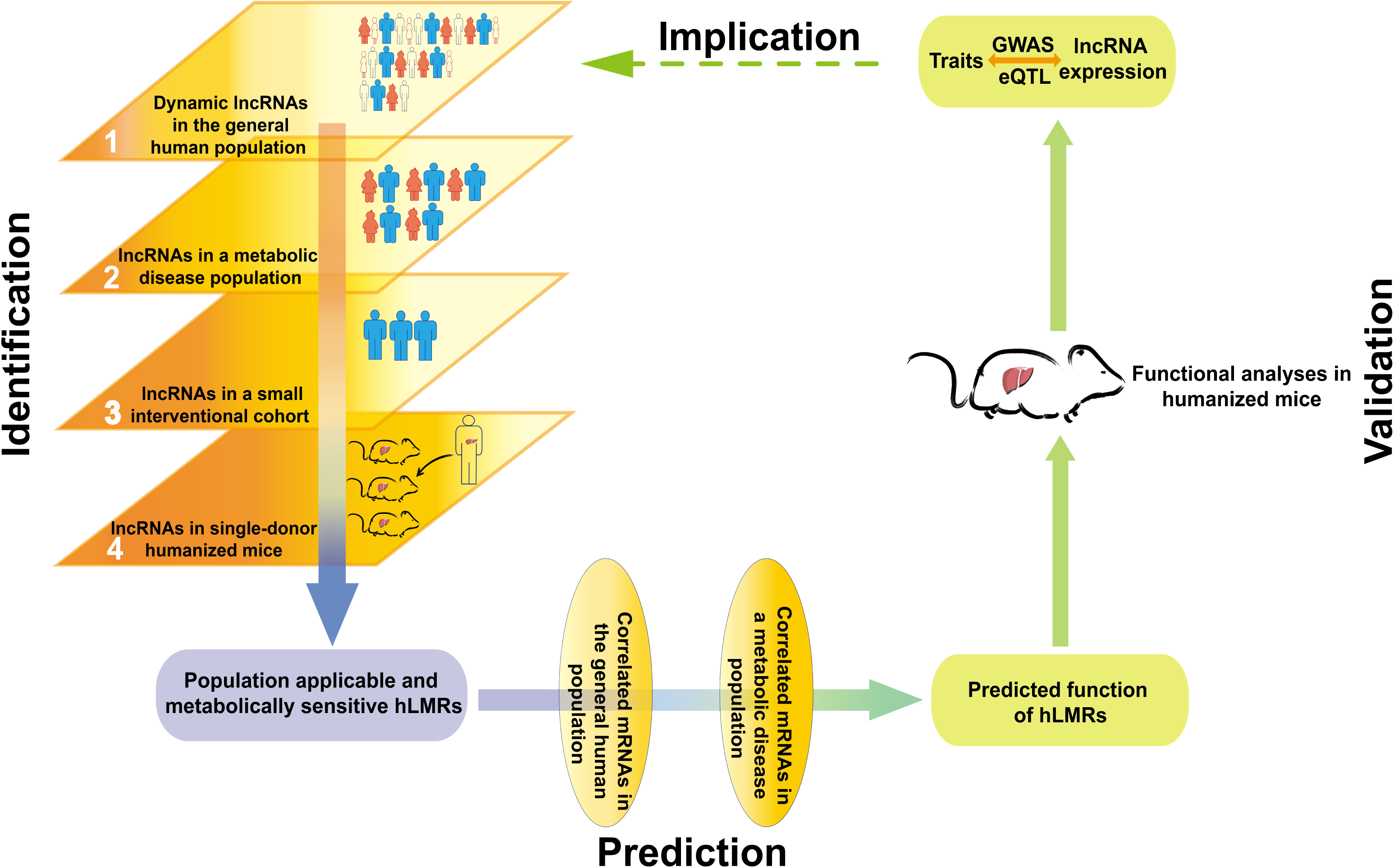
A roadmap for defining human lncRNA metabolic regulators (hLMRs) An integrative roadmap illustrates the steps to identify population applicable, metabolically sensitive and disease-relevant hLMRs by step-wise selections of human lncRNAs in the general population, a metabolic disease population, a small disease cohort and humanized mice (Identification), to infer their function based on their correlation with mRNAs in multiple populations (Prediction), to validate their function in humanized mice (Validation) and finally to explore their relevance to human diseases (Implication).

**Figure 2.**
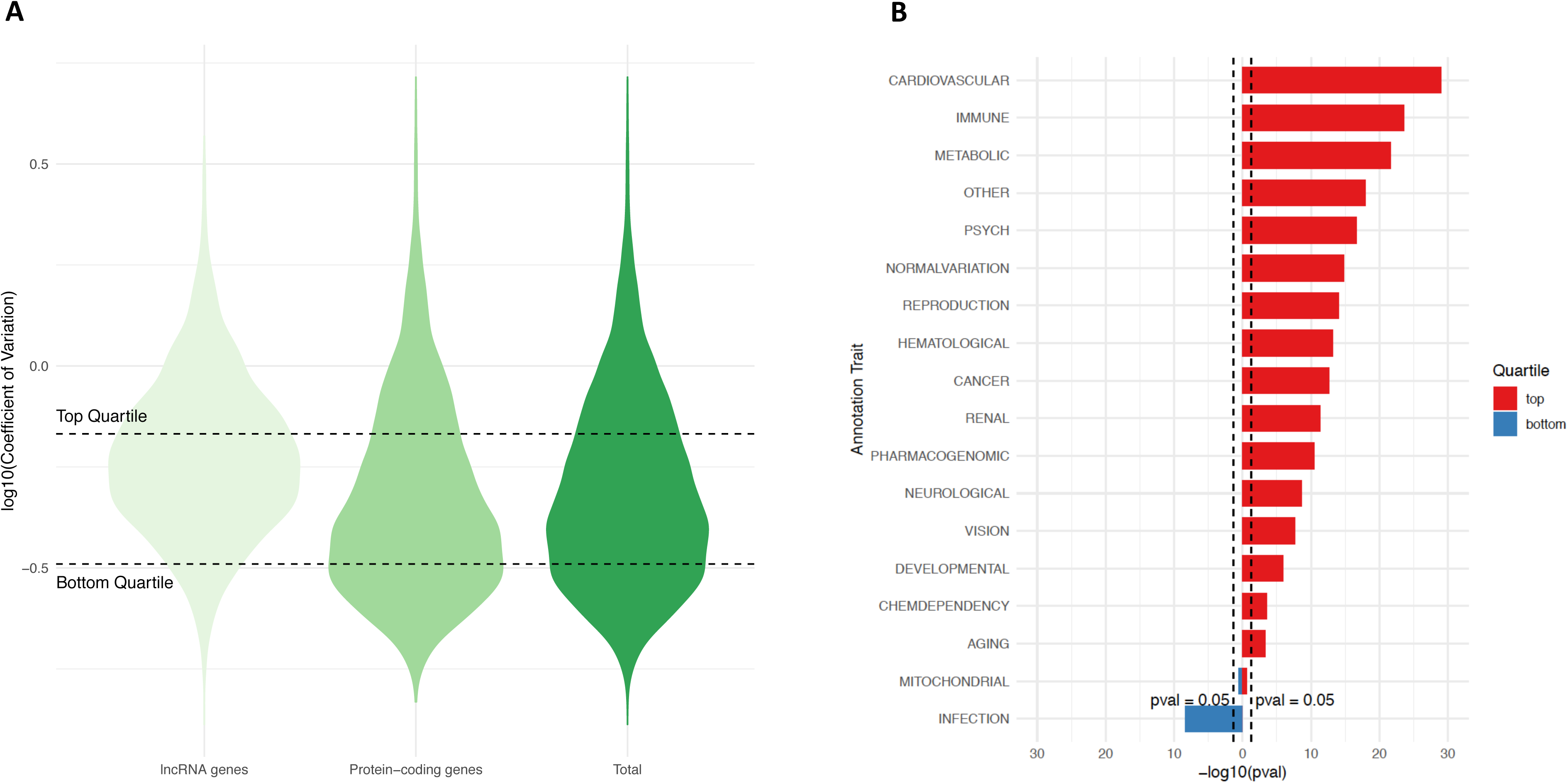
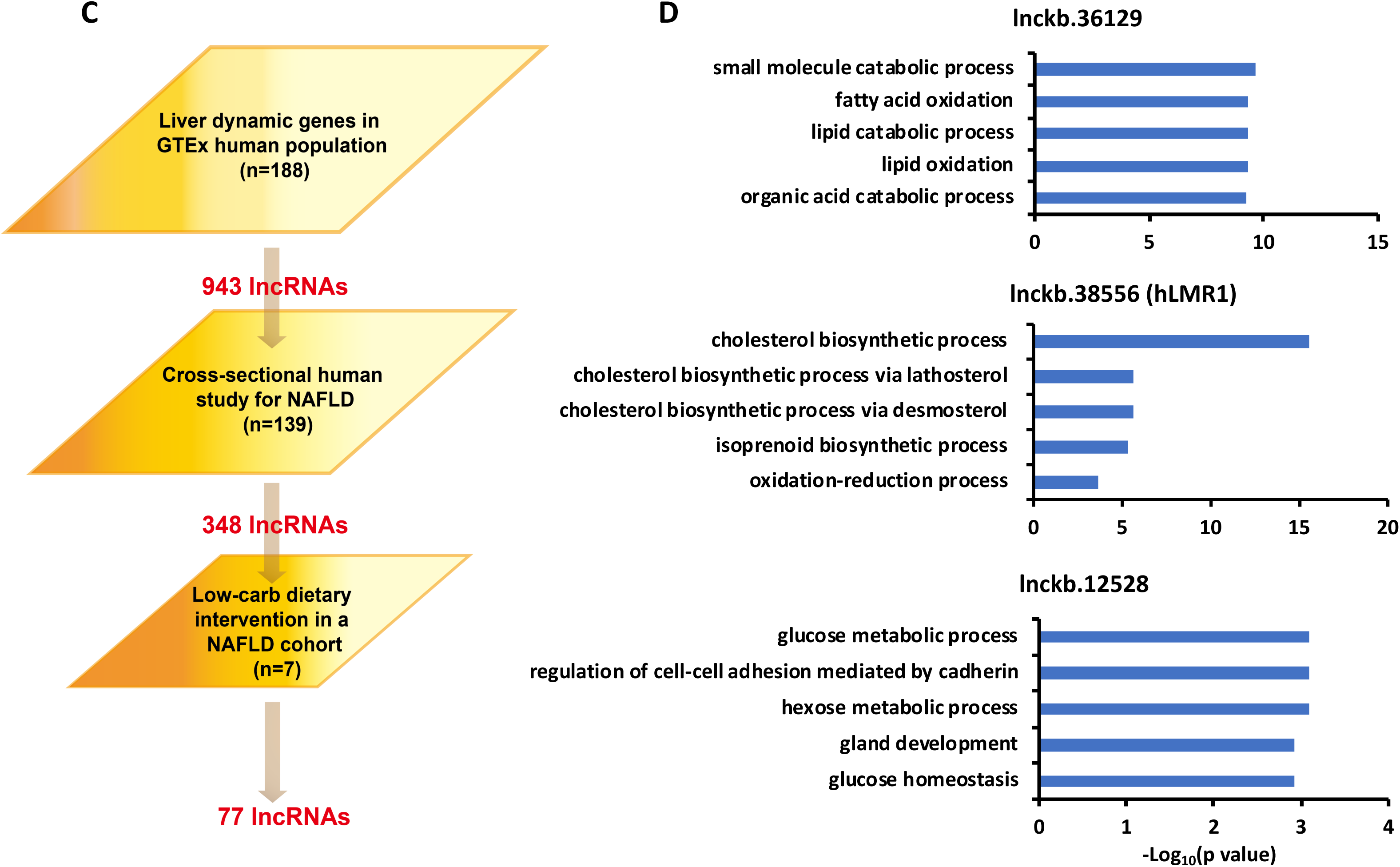
Identification of human lncRNA metabolic regulators (hLMRs) (A) Violin plots of Coefficient of Variation of expressed genes (n=16906) in GTEx liver dataset (n=188; scaled by area). Coefficient of Variation was log_10_ normalized. (B) Enrichment of the disease classes assigned to the protein-coding genes in bottom quartile (left) and top quartile (right) dynamically expressed in GTEx liver dataset. (C) The process of identification of hLMRs by multilevel analysis of different types while relevant human data. The numbers of lncRNAs resulted from each additional step are marked in red. NAFLD, Non-Alcoholic Fatty Liver Disease. (D) GO term analyses using the protein-coding genes persistently correlated with hLMRs in independent human populations. Only the top five GO terms of each hLMR were presented. See also Tables S1 to S4

To determine the responsiveness of these dynamic lncRNAs to a specific metabolic disease condition, we examined their regulation in non-alcoholic fatty liver disease (NAFLD). NAFLD is a metabolic disorder of high prevalence and is known to cause global changes in gene expression and metabolism in the liver. Specifically, we analyzed a large RNA-seq dataset composed of liver samples (total 139 samples) from a cross-sectional study of NAFLD^23^ (Figure 2C and Table S2) and found that 348 of 943 (∼37%) dynamically expressed lncRNAs are differentially expressed in the NAFLD populations, further supporting the notion that highly dynamic lncRNAs may have direct relevance to metabolic disease (Figure 1, Identification steps 1-2).

Although the 348 lncRNAs have potential implications for both general and disease populations, their observed changes in expression levels could be affected by genetic heterogeneity inherent to population-based human studies. To minimize such effects and further enrich lncRNAs whose expression levels are indeed metabolically sensitive, we next analyzed a liver RNA-seq dataset generated from liver biopsies of seven obese NAFLD patients before and after a short-term intervention with a low-carb diet^24^. Low-carb diet is known to cause rapid and robust reductions of liver fat as well as changes in the expression of genes in multiple metabolic pathways^24^. Out of the 348 lncRNAs, 77 are also regulated by the low-carb dietary interventions in this small cohort (Table S3 and S4 and Figure 1, Identification step 3). Thus, by integrating gene expression dynamics in the general population and gene regulation in both observational and interventional human studies related to NAFLD, we have effectively identified a group of population applicable, metabolically sensitive and disease-relevant human lncRNAs, which we refer to as hLMRs (human lncRNA metabolic regulators) (Figure 1, Identification steps 1-3).

As presently we cannot deduce lncRNA function from their sequence, it has become a common practice to predict a lncRNA’s function based on its correlated protein-coding genes in a large transcriptome dataset^12^. On many occasions, however, the number of such correlated genes could be very large, and the predicted functions could be very broad. Additionally, the correlated genes often vary significantly in distinct populations. To overcome these limitations, we extract genes persistently correlated with a lncRNA in multiple independent datasets, which we postulated could remove some protein-coding genes that are spuriously co-regulated with a lncRNA and enrich the lncRNA’s specifically correlated genes. Specifically, we intersected protein-coding genes correlated with a lncRNA in both general GTEx population and in samples from the NAFLD study described above. As a result, we identified a concise list of correlated genes of each hLMR in a metabolic disease-relevant setting. The Gene Ontology (GO) analysis using these lists indicates that our identified hLMRs may function in diverse metabolic pathways such as fatty acid oxidation, cholesterol biosynthetic process and glucose metabolism (Figure 2D and Table S4).

### Metabolic Regulation of hLMRs in humanized mice

We have undertaken considerable efforts to filter out confounding variables to identify metabolic sensitive hLMRs and predict their functions. We next asked if we can further investigate their metabolic responses under a physiologically relevant and well-controlled experimental condition. Indeed, we have recently found that a liver-specific humanized mouse model, which is produced by human hepatocytes from a single donor and are kept under defined environment in an animal facility^25^, is suitable for studying the regulation of human-specific lncRNAs. We thus performed RNA-seq analysis to identify differentially expressed human genes in the liver-specific humanized mice subjected to a fasting-refeeding regime, which involves the two extreme ends of caloric cycles and is known to regulate nearly all key metabolic genes in vivo (Table S5). As expected, while the expression of protein-coding genes involved in fatty acid oxidation and gluconeogenesis are upregulated by fasting and downregulated by refeeding, genes in the lipogenesis pathway show the opposite pattern (Figure 3A), supporting the proper response of human genes to nutrient and hormonal levels in the humanized liver. Furthermore, we noticed that a significant portion of differentially expressed human genes during fasting and refeeding overlap with those in the NAFLD and the low-carb dietary intervention analysis (Figure 3B). These results further support the humanized liver maintains an appropriate gene expression response to metabolic milieu as the human liver dose. Finally, we found 20 out of 77 of the liver hLMRs that we have identified are regulated by feeding cycles in the humanized mice (Figure 3C). Indeed, the specific regulations of these 20 hLMRs are largely in line with their predicted function. For example, lnckb.38556, which is downregulated by fasting and recovered by refeeding, is predicted to function in biosynthesis of cholesterol.

**Figure 3.**
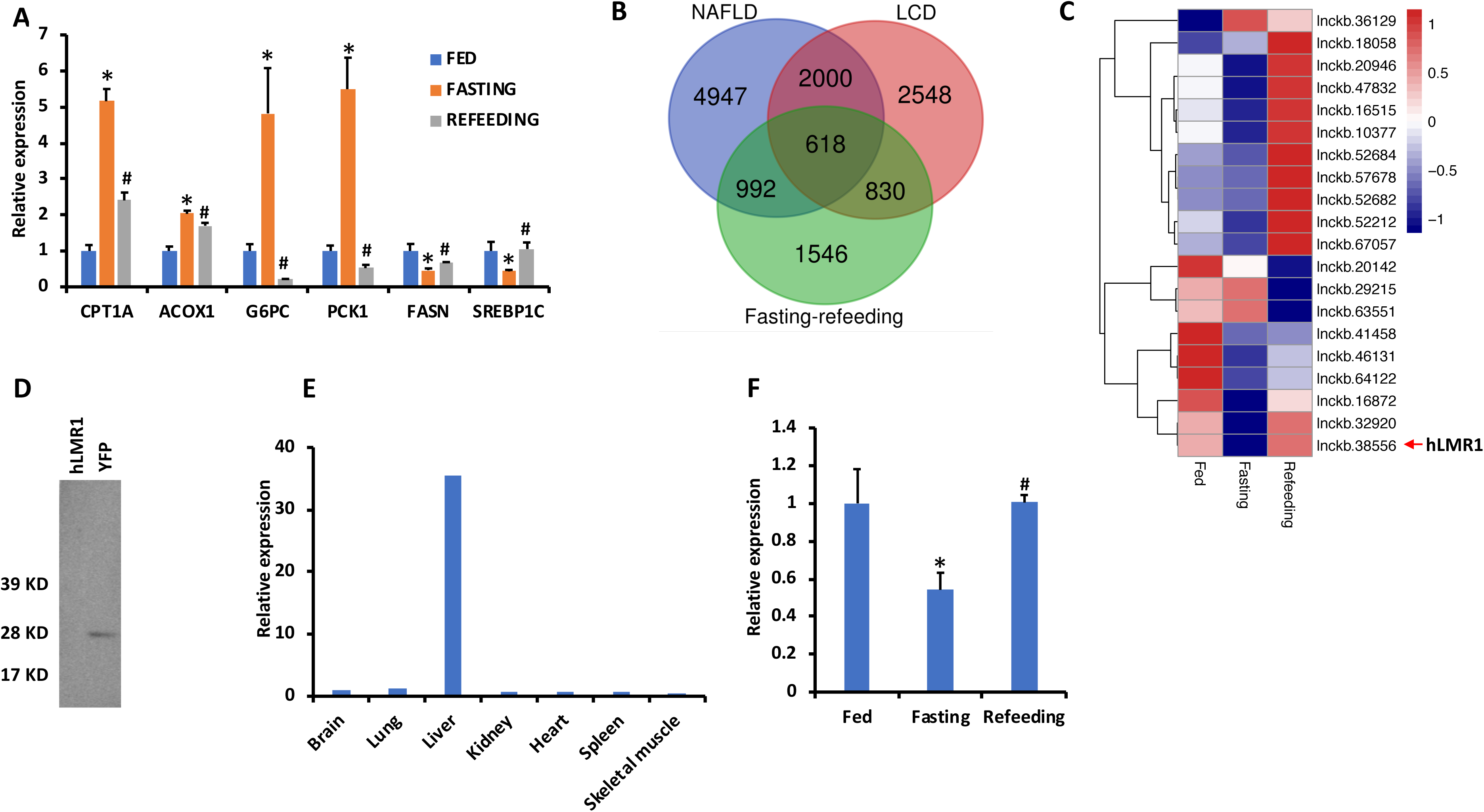
Metabolic Regulation of hLMRs in humanized mice. (A) Gene expression in the livers from the humanized mice in response to fasting and refeeding (Fed, n=4; Fasting, n=5; Refeeding, n=5), error bars represent SEM, * p<0.05 for Fasting vs Fed, # p<0.05 for Refeeding vs Fasting. (B) Venn diagram for the intersection between metabolic responsive genes in humanized mice, the differentially expressed genes in non-alcoholic fatty liver disease population (NAFLD) and the regulated genes by low-carbohydrate dietary intervention (LCD). (C) Heatmap of the relative mean expression levels for the hLMRs in the livers from fed, fasted and refed humanized mice. hLMR1 was pointed out by a red arrow. (D) In vitro translation analysis for hLMR1 and the coding sequence of YFP. (E) Relative expression levels of hLMR1 across different human tissues. Data were normalized to the expression level in brain (set as 1). (F) Expression of hLMR1 in the liver of humanized mice in response to fasting and refeeding (Fed, n=4; Fasting, n=5; Refeeding, n=5), error bars represent SEM, * p<0.05 for Fasting vs Fed, # p<0.05 for Refeeding vs Fasting. See also Tables S5.

Taken together, by performing stepwise selections of lncRNAs from multiple datasets representing the general population, disease populations, interventional studies, and well-controlled experiments in humanized mice, we have established a list of potential hLMRs that are population applicable, metabolically sensitive and disease-relevant (Figure 1, Identification steps 1-4). Furthermore, by extracting protein-coding genes that are persistently correlated with lncRNAs in independent populations, we were able to generate a concise list of genes which could be utilized to infer the function of each hepatic hLMR for downstream analysis (Figure 1, Prediction steps).

### Regulation of hepatic cholesterol biosynthetic pathway by hLMR1 in the humanized liver

Among the 20 liver hLMRs, we used lnckb.38556, which we refer to as hLMR1, as an example to experimentally validate our selection and prediction process. hLMR1 is a five-exon intergenic lncRNA located in chromosome 3 of human genome, and its annotated full-length transcript is 2kb (Figure S1A). No homolog of hLMR1 can be identified in mice by a BLAST search, suggesting it is a non-conserved human lncRNA. In addition to bioinformatic prediction, an in vitro translation assay further supports that hLMR1 is a non-coding gene (Figures 3D and S1B). Our quantitative Real-time PCR (qPCR) results using human tissue cDNA panels showed that hLMR1 is exclusively expressed in human liver tissue (Figure 3E) with an estimated copy number per cell of 56. Subcellular fractionation analysis using humanized liver tissues found that hLMR1 is distributed in both of cytoplasm and nucleus with more in the nuclear fraction (Figure S1C). RNA-seq analysis of humanized livers showed that hLMR1 is down regulated during fasting and recovered upon refeeding (Figure 3C), which can be further verified by qPCR (Figure 3F). Furthermore, GO term analysis using the list of genes persistently correlated with hLMR1 indicates that hLMR1 may function in the cholesterol biosynthetic processes (Figure 2D), which is consistent with the observation that hLMR1 is induced by refeeding.

To directly test the predicted role of hLMR1 in cholesterol biosynthetic pathway, we first screened for shRNAs that could efficiently block the expression of hLMR1 and then used adenoviruses expressing the selected shRNA to specifically reduce the expression levels of hLMR1 in the livers of humanized mice. As shown in Figure 4A, this strategy successfully depleted the expression of hLMR1 by more than 70%. Remarkably, among the six crucial genes in the cholesterol biosynthetic pathway whose expressions are correlated with hLMR1 in our GO term analysis, four genes including SC5D, FDPS, LSS, and HMGCS1 showed decreased expression by more than 50% upon depletion of hLMR1 in the humanized livers (Figure 4A). This result thus supports that hLMR1 positively regulates the cholesterol biosynthetic pathway as predicted. We noticed that depletion of hLMR1 had no effect on the expression of PAQR9, the close neighbor gene of hLMR1 (Figure 4A), indicating a trans acting mechanism of hLMR1.

**Figure 4.**
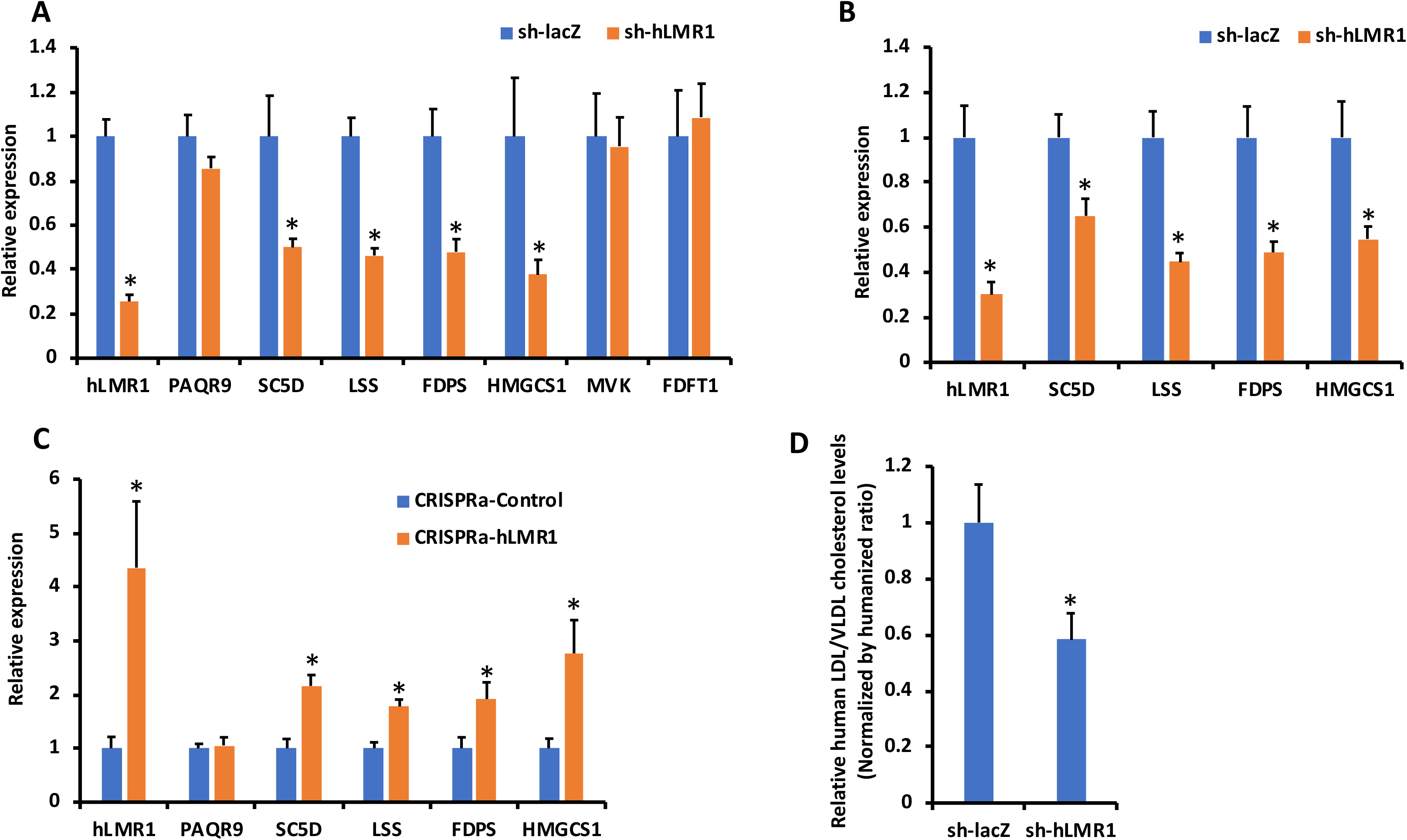
Regulation of cholesterol metabolism by hLMR1 in humanized mice. (A) Gene expression in humanized mice receiving adenovirus for control (sh-lacZ, n=5) or knocking down of hLMR1 (sh-hLMR1, n=6). (B) Gene expression in humanized mice (2^nd^ donor) receiving adenovirus for control (sh-lacZ, n=4) or knocking down of hLMR1 (sh-hLMR1, n=4). (C) Gene expression in humanized mice receiving adenovirus for CRISPRa-Control (n=4) or CRISPRa-hLMR1 (n=4). (D) Relative cholesterol levels in human ApoB-containing lipoproteins purified from the plasma of humanized mice receiving adenovirus for control (sh-lacZ, n=7) or knocking down of hLMR1 (sh-hLMR1, n=10). Error bars represent SEM, * p<0.05. See also Figure S1.

As our pipeline is designed to identify human lncRNA metabolic regulators implicated in general population, we next test if the regulatory effects of hLMR1 could be observed in a different genetic background. Knockdown experiment as described above was hence performed in humanized mice prepared with hepatocytes from a second independent and ethnically different donor. As shown in Figure 4B, with a similar knockdown efficiency of hLMR1 in these mice, we found significant downregulation of SC5D, LSS, FDPS and HMGCS1, which is consistent with the result we observed in mice produced with the first donor. Taken together, our results support that hLMR1 is critical to maintain the expression of cholesterol biosynthetic genes in human populations.

To further study the regulatory effects of hLMR1, we next asked if overexpression of hLMR1 could promote the expression of genes in cholesterol biosynthetic pathway. To this end, we took advantage of the CRISPR activation (CRISPRa) tool to enhance the expression of endogenous hLMR1 specifically in human hepatocytes of the humanized liver. As shown in Figure 4C, adenovirus-mediated expression of CRISPRa targeting hLMR1 in humanized mice induced the expression level of hLMR1 by 4 folds, while the expression of PAQR9 was not affected. Corroborating the result of our knockdown experiments, we found that induction of hLMR1 by CRISPRa resulted in significant upregulation of SC5D, FDPS, LSS, and HMGCS1. This data further support that hLMR1 is not only necessary for maintaining the expression of cholesterol biosynthetic genes, but also sufficient to promote their expressions.

The regulatory effects of hLMR1 on cholesterol biosynthetic genes encouraged us to examine if hLMR1 could modulate cholesterol levels in the humanized mice. As the humanized livers in these mice are chimeric and it is technically challenging to ascertain the specific impact of human hepatocytes. We thus used an immunoaffinity approach to specifically isolate human ApoB-containing lipoproteins from the plasma of humanized mice (Figure S1D and Methods). Using this method, we found that depletion of hLMR1 in humanized livers led to 40% decrease in human LDL and VLDL cholesterol levels compared with control humanized mice (Figure 4D). Taken together, our bioinformatic analyses using large scale human data as well as functional analyses in humanized livers support that a non-conserved human lncRNA, hLMR1, is a crucial regulator of hepatic cholesterol biosynthetic pathway and could play a critical role in maintaining cholesterol homeostasis in human.

### hLMR1 coordinates PTBP1 to promote the transcription of cholesterol biosynthetic genes

To explore the molecular mechanism mediating the regulatory effects of hLMR1, we first performed RNA poly II ChIP analysis to determine the transcriptional activities of the hLMR1 target genes in the humanized livers. As shown in Figure 5A, humanized livers with depletion of hLMR1 showed significant lower enrichment of RNA poly II on the transcription start site (TSS) of human SC5D, FDPS, LSS, and HMGCS1. This data suggest that hLMR1 regulates the expression of cholesterol biosynthetic genes by promoting their transcription, in line with the relative enrichment of hLMR1 in nuclear fraction (Figure S1C). To further explore how hLMR1 regulates the transcription of its target genes, we next performed RNA pulldown combined with mass spectrometry analysis to identify proteins that interact with hLMR1. This strategy successfully identified PTBP1, an RNA-binding protein regulates almost all steps of RNA metabolism, as a hLMR1-binding partner. We confirmed the mass-spectrometry results by western blot analysis (Figure 5B), and then a RIP analysis using PTBP1 antibody in liver tissue lysate of humanized mice was performed. As shown in Figure 5B, hLMR1 is enriched by more than 50 folds in PTBP1 immunoprecipitate as compared to IgG, further supporting the interaction between hLMR1 and PTBP1. To directly test if PTBP1 is involved in hLMR1 mediated transcriptional regulation of cholesterol biosynthetic genes, we overexpressed hLMR1 in combination with PTBP1 to determine their effects on a human HMGCS1 promoter-driven luciferase reporter. As shown in Figure 5C, expression of either hLMR1 or PTBP1 could increase the HMGCS1 promoter-driven luciferase activity, and simultaneous expression of both of them showed a synergistic enhancing effect. This result suggests that PTBP1 is a positive transcriptional regulator of cholesterol biosynthetic genes, and hLMR1 likely function through facilitating the recruitment of PTBP1 to the promoters of its target genes. To experimentally test this, we performed PTBP1 ChIP analyses in the humanized liver and determined PTBP1 enrichment on the promoters of hLMR1 target genes. As shown in Figure 5D, PTBP1 exhibited robust enrichment on the promoters of human SC5D, FDPS, LSS, and HMGCS1, and its binding to these promoters was diminished by depletion of hLMR1 in humanized livers. Taken together, our results support that hLMR1 recruits PTBP1 to the promoters of genes in the cholesterol biosynthesis pathway to activate their transcription.

**Figure 5.**
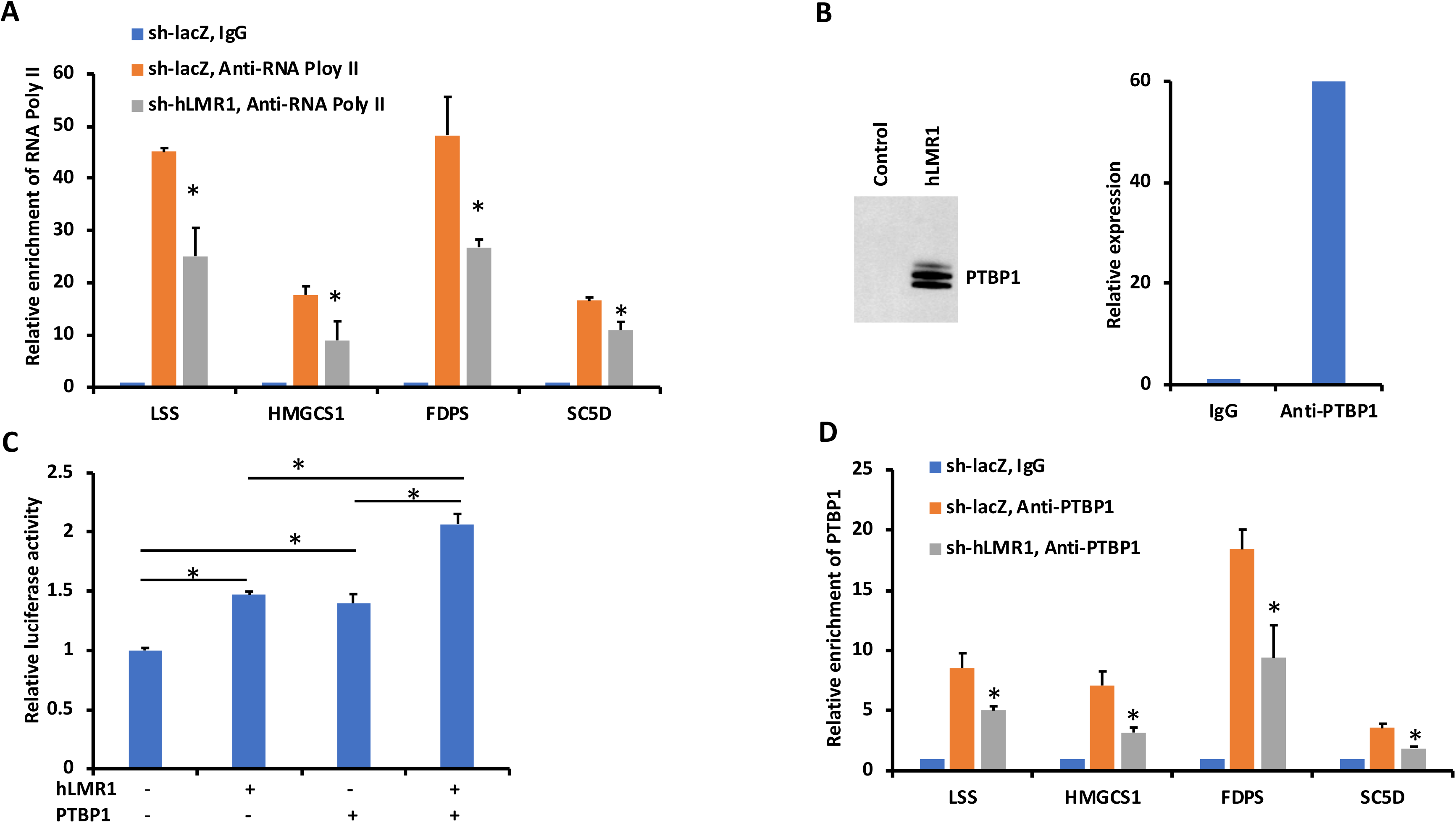
hLMR1 coordinates PTBP1 to promote the transcription of cholesterol biosynthetic genes. (A) RNA ploy II ChIP analyses in liver tissues of humanized mice receiving adenovirus for control (sh-lacZ, n=3) or knocking down of hLMR1 (sh-hLMR1, n=3). (B) Left: western blot analysis of PTBP1 in hLMR1 pulldown, right: expression of hLMR1 in PTBP1 RIP (RNA immunoprecipitation) in humanized liver. (C) HMGCS1 promoter-driven luciferase reporter assay in 293A cells (n=3 for each group). Data are representative results of three independent experiments. (D) PTBP1 ChIP analyses in liver tissues of humanized mice receiving adenovirus for control (sh-lacZ, n=3) or knocking down of hLMR1 (sh-hLMR1, n=3). Error bars represent SEM, * p<0.05.

### Ectopic expression of hLMR1 in regular mice promotes biosynthesis of cholesterol

To further study hLMR1-PTBP1 mediated regulation of cholesterol metabolism, we asked if the function of human PTBP1 is conserved in mouse and as such, ectopic expression of hLMR1 in regular mice could promote biosynthesis of cholesterol. To test these possibilities, we used adenovirus mediated shRNAs to knock down Ptbp1 in regular mice and then examined the expressions of cholesterol biosynthetic genes. As shown in Figure 6A, when more than 70% Ptbp1 was depleted, we observed decreased expression of Sc5d, Fdps, Lss and Hmgcs1. This result supports that mouse Ptbp1 also positively regulates the expression of cholesterol biosynthetic genes which allows us to test the effect of ectopic expression of hLMR1 in regular mice. We thus cloned the full-length cDNA of hLMR1 in an adenoviral vector and delivered the packed adenoviruses into regular mice. This strategy successfully expressed hLMR1 in mouse livers to the level that is comparable to that in human hepatocytes. As shown in Figure 6B, we found expression of hLMR1 resulted in marked up-regulation of Sc5d, Fdps, Lss and Hmgcs1 in the liver of regular mice. Furthermore, both plasma and hepatic cholesterol levels were increased upon expression of hLMR1 (Figure 6C). These data, together with our results in humanized mice, support the crucial role of hLMR1-PTBP1 complex in cholesterol metabolism.

**Figure 6.**
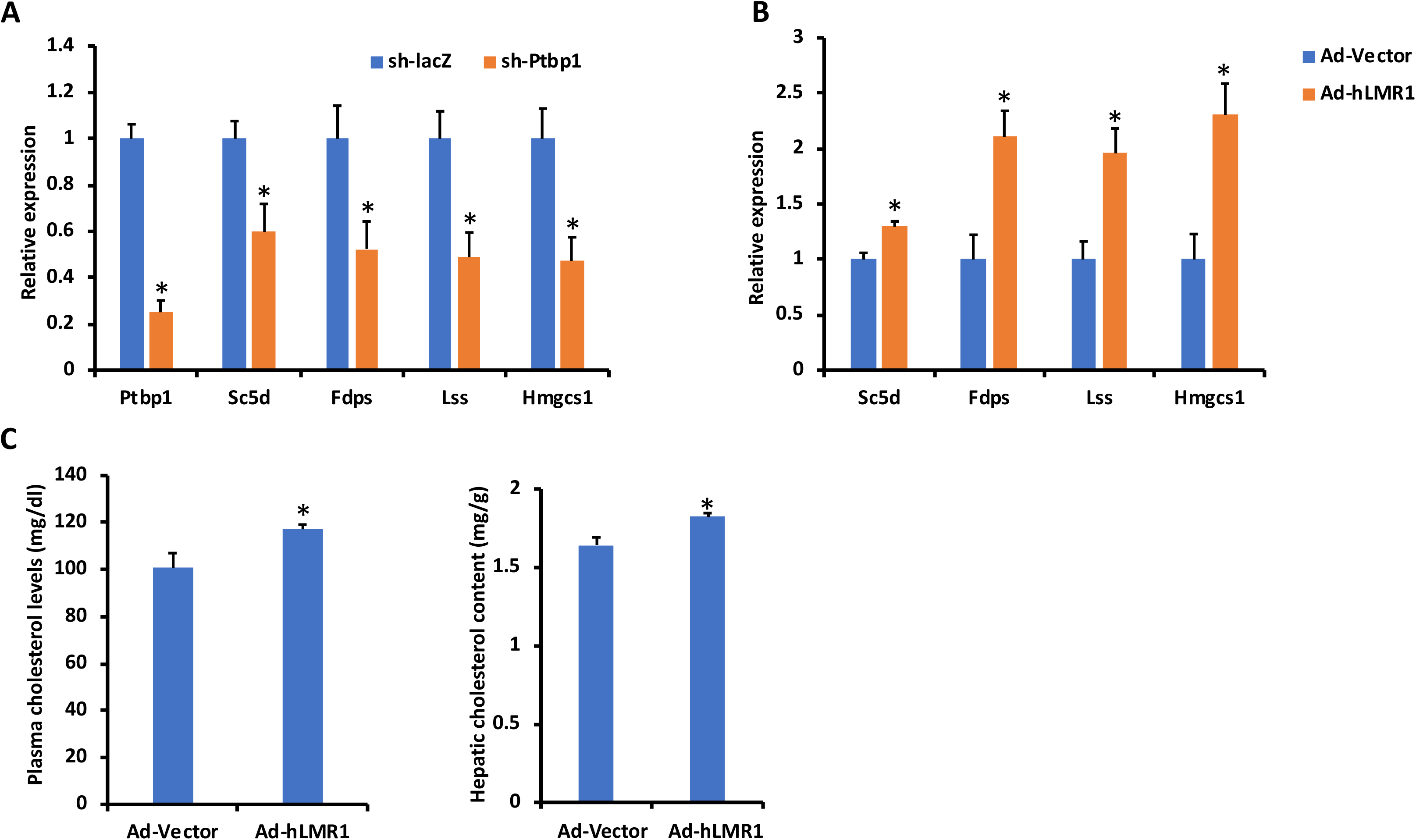
Ectopic expression of hLMR1 in regular mice promotes biosynthesis of cholesterol. (A) Gene expression in the liver of regular mice receiving lacZ shRNA (sh-lacZ, n=9), or shRNA for Ptbp1 (sh-Ptbp1, n=7). (B) Gene expression in the liver of regular mice receiving adenovirus for control (Ad-Vector, n=5) or expression of hLMR1 (Ad-hLMR1, n=6). (C) Plasma (left) and liver (right) cholesterol levels in regular mice receiving adenovirus for control (Ad-Vector, n=9) or expressing of hLMR1 (Ad-hLMR1, n=9). Error bars represent SEM, * p<0.05.

### The hepatic expression of hLMR1 is associated with cholesterol levels in human population

To further explore the implication of hLMR1 mediated regulation of cholesterol metabolism in the human population, we used eQTL-GWAS integrative analysis to determine the association between the hepatic expression of hLMR1 and the lipid levels in the general population (Cookson et al., 2009). As shown in Figure 7, encouragingly, we found several cis eQTLs of hLMR1 overlap with GWAS loci for total cholesterol levels. An SMR (Summary-data-based Mendelian Randomization) analysis, which tests if the effect size of a SNP on the phenotype is mediated by gene expression using data from GWAS and eQTL studies (Zhu et al., 2016), was then performed to further determine whether the overlapped eQTL/GWAS loci are functionally related. The analysis passed both SMR and HEIDI (heterogeneity in dependent instruments) tests (pSMR<0.05 and pHEIDI>0.05, see Methods), supporting the hepatic expression of hLMR1 might contribute to the regulation of cholesterol levels in human.

**Figure 7.**
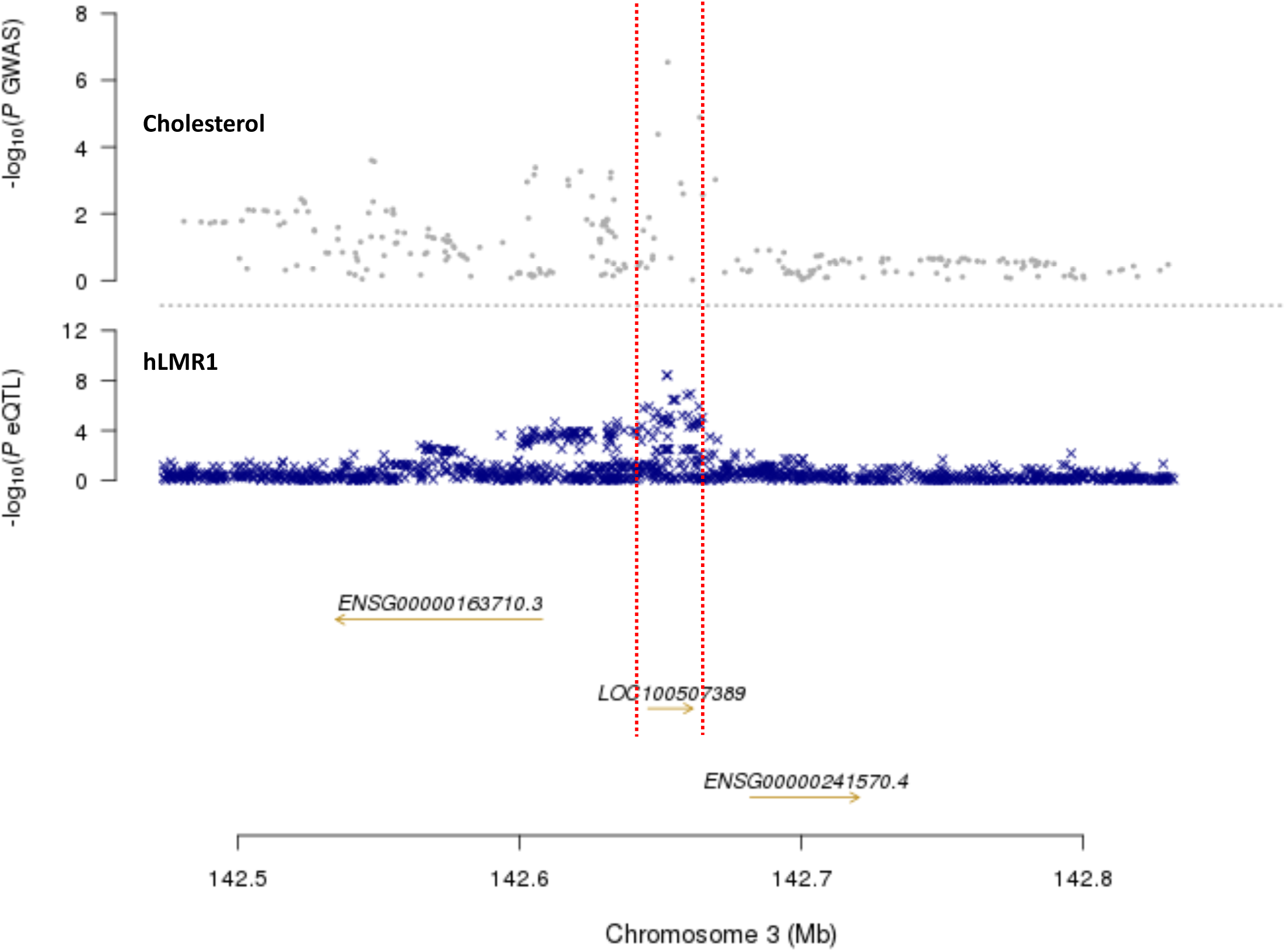
The illustration of the overlap between eQTLs of hLMR1 and GWAS loci for lipid trait.

## Discussion

Human lncRNAs constitute a significant portion of the human transcriptome and have been shown to play a critical role in diverse biological processes. Despite this, the role of human lncRNAs in systemic energy metabolism is poorly understood, in part, because of the challenging nature of identifying hLMRs and defining their metabolic function in a physiologically relevant context. In this work, we established an integrated bioinformatic and experimental pipeline to identify hLMRs based on their regulatory information in general population, patients with metabolic disease, and a humanized mouse model (Figure 1). We also adapted an improved approach to predict the function of hLMRs. Finally, we provided a proof of principle example that the metabolic function of human lncRNAs can be successfully defined in the humanized mouse model, where we validated that a non-conserved human lncRNA exhibited its predicted function in lipid metabolism and confirmed it as a legitimate hLMR.

Due to the limitations in using sequence features to identify hLMRs, examining the change in expression of human lncRNAs in response to metabolic conditions is a major tool to establish their potential role in metabolism. Compared to data generated in inbred mice under well-defined conditions, however, human gene expression data are intrinsically very noisy due to multiple confounding factors, particularly the diverse genetic background and environmental factors. On the other hand, steps to remove the impact of genetic and other confounding factors have the risk of restricting the significance of the findings to a specific subpopulation. To address these challenges, our work has established a platform to enhance the identification of hLMRs by implementing several steps to better retrieve specific signals against the background noise in human data and at the same time retain the significance of identified lncRNAs to the general population: 1) we started by selecting highly dynamic lncRNAs in the general human population by utilizing a most updated and comprehensive lncRNA annotation we have recently generated to capture the hLMRs in key metabolic organs. Protein-coding genes yielded by this analysis are enriched in disease-relevant states, suggesting that these highly dynamic lncRNAs may also be relevant to metabolic disease processes; 2) we next selected lncRNAs that are regulated by a common metabolic disease condition, and lncRNAs identified in this study are expected to be both disease-relevant and represented in the general population; 3) we subsequently used interventional studies, although usually in a relative small cohort, to further eliminate the influence of genetic and environmental effects and to enrich lncRNAs that are specifically regulated by metabolic milieu; 4) we used well-controlled experiments in a humanized mouse model to complement the regulatory information derived from human liver samples, further reducing the influence of any confounding factors while keeping the direct implication in humans. In theory, we could potentially select functional lncRNA candidates from the regulated lncRNAs in humanized mice but we are mindful of the potential artifacts from a model system and believe that the combination of regulatory information in both human and humanized system yields the most representative candidates; 5) finally, we improved the approach to generate a short list of co-expressed coding genes for functional prediction of each liver hLMR, which we have subsequently shown to be vital in designing hypothesis-driven experiments. Furthermore, our pipeline is entirely scalable and can readily integrate any new RNA-seq data of human metabolic organs under any additional metabolic conditions. These data are expected to become available en masse with the wide-spread use of RNA-seq technology. LncRNA research is currently at an interesting juncture, with the focus of the field quickly transitioning from lncRNA annotation to their functional definition. Our analytical/experimental pipeline, and the multiple selective lists of potential hLMRs, are aimed at providing a timely resource for more metabolism labs to further characterize hLMRs and understand their overall impact on metabolic homeostasis, which may yield novel insights into the regulation of human metabolism and fill gaps in our understanding of the pathophysiology of metabolic diseases.

Our extensive analyses have generated a very selective list of human lncRNAs, with their potential functions, which would significantly facilitate the subsequent experimental validation of the specific functions of hLMRs. To study the functions of hLMRs in a physiologically relevant context, we used a liver-specific humanized mouse model. This model has re-established the bi-directional information flow between humans and experimental mice, which has broken down for lncRNAs due to their low conservation. As shown by the example of hLMR1, this model is crucial for the definitive validation of the physiological importance of any putative hLMRs, which are mostly non-conserved. Of note, some non-conserved lncRNAs can indeed be studied in cultured cells as shown in two recent reports^26, 27^. Clearly, these studies can be significantly enhanced if the same targets can be further investigated in an in vivo model such as the humanized mice in the current study. Of course, the humanized mouse model has its limitations. Due to the size of the mouse liver and immunodeficiency of the mouse strain, it is not expected to have the capacity to recapitulate all aspects of human liver biology. Nonetheless, to our knowledge, it currently represents a system that matches the human liver as closely as currently possible. Of note, we have also demonstrated that expression of human lncRNAs in regular mice is a useful alternative to the humanized mice. This approach may mitigate some of the limitations of studying human lncRNAs in mice. However, the human lncRNA expression in mice cannot replace the utility of the humanized mice in which the lost-of-function of human lncRNAs provides a nearly definitive confirmation of their relevance to human physiology.

LncRNAs are the largest and probably also the least conserved transcript class in the human genome^15^. Mounting evidence of functional human lncRNAs in diverse biological processes suggests that they may significantly contribute to interspecies differences, particularly among mammals. Although it is still speculative at this stage, it is plausible that the functional impacts of many non-conserved human lncRNAs that are unapproachable/inaccessible in animal studies might be partly responsible for the very low translation rate of mouse-based studies to human therapies. Still, the reasons for our current limited understanding of lncRNA function clearly go beyond their recent discovery, and the combination of low conservation of lncRNAs restricting regulatory information of human lncRNAs to human data and the lack of technological approaches to specifically process human data to accurately define lncRNA regulation are likely major contributing factors. Thus, our integrated analysis in human and humanized mouse model have enabled human lncRNA candidate selection, functional inference, and validation of hLMR function. We propose that this approach will open up a novel avenue to more rapidly identify and define the functional significance of human lncRNAs in the pathophysiology of metabolic disorders. With the growing importance of human lncRNAs in biology and physiology, defining hLMRs could drive profound changes in the way we study energy metabolism experimentally and understand metabolic disease conceptually. We hope that our resource and pipeline will help chart the way forward.

## Methods

### Bioinformatics Analysis

#### Analysis pipeline for human RNA-seq data

Fastq read files were cleaned using TrimGalore. Reads were then aligned using HISAT2 to an index created using the GRCh38 genome and the lncRNAKB annotation. Aligned .sam files were then compressed into .bam files and sorted using Sambamba sort. Feature Counts from the subread package was used to count reads/fragments aligned to genes (at the exon feature level). Samples with less than 1,000,000 reads aligned were removed. Individual count files were merged into a single count file for each dataset using a python script. In each dataset, a threshold of >1 cpm in at least 50% of samples was applied before further analysis. The RNA-seq datasets for cross-sectional human studies are retrieved from BioProject PRJNA512027 for the 139 liver samples from the non-alcoholic fatty liver disease population (NAFLD). The RNA-seq datasets for interventional human studies are retrieved from BioProject PRJNA420975 for the 7 paired liver samples of low-carb dietary intervention in NAFLD at NCBI database.

#### Analysis pipeline for humanized mice RNA-seq data

RNA-seq data have been deposited in NCBI Gene Expression Omnibus (GEO) (https://www.ncbi.nlm.nih.gov/geo/query/acc.cgi?&acc=GSE130525, reviewer token: spsnkqcadtupvuf). A combined human and mouse annotation was created by combining the lncRNAKB gtf annotation with the RefSeq GCF000001635.26 gtf annotation. A combined human and mouse genome was created by combining the GRCh38.p12 Genome sequence, primary assembly obtained from the Gencode website and the GRCm38.p6 genomic sequence obtained from the refseq ftp site. RNA-seq cleaning, alignment, sorting, quantification and filtering were conducted in the same way as Human RNA-seq analysis using this index and genome file. Human gene counts were extracted from the resulting count file by removing reads mapped to mouse contigs. A >1 cpm in 50% of samples cutoff was applied separately to human gene counts and mouse gene counts.

#### Differential expression analysis and PCA

A combined raw count file generated by the Subread featureCounts tool for each dataset was imported into R. Technical replicates were combined using the collapseReplicates function from the DESeq2 package. The variance stabilizing transform from the DESeq2 package was applied to the count data from before and after combining technical replicates (if there were technical replicates) before conducting principal component analysis (PCA). The top two PCs were graphed to visualize clustering between experimental groups. DESeq2 was used with non-normalized count data to find differentially expressed genes between experimental groups. Covariates were controlled for by adding them to experimental design if available. A cutoff of logFC>0.4 and q<0.05 is used for differential expression for all the liver samples.

#### Gene Variability Analysis

The gene expression profiles of human liver from GTEx v7 includes samples from caucasian, asian, and african ancestry (primarily caucasian). An expression cut off of >1 cpm in 50% samples was applied in each tissue to reduce mapped genes to 16906 expressed genes including 2665 lncRNA genes in the liver. For each of the expressed genes, we quantified expression variability by calculating its coefficient of variation η across all the available samples in each tissue. These were subsequently ranked and split into quartiles. Diseases category analysis was performed by using DAVID gene functional annotation tool.

#### Gene correlation analysis

Count data for each data set was Variance Stabilizing Transform (VST) normalized using the DESeq2 R package. After VST normalization, 10 hidden technical factors were calculated per data set using the Probabilistic Estimation of Expression Residuals (PEER) software package and were used in linear regression as covariates to correct VST normalized expression data. The mean expression of each gene was added back to the expression residual. The correlation between lncRNAs and protein coding genes was analyzed by Pearson’s method using the normalized data in GTEx human population and metabolic disease-relevant population. Significantly correlated protein coding genes with p<0.05 in the liver were used for further Gene Ontology (GO) analysis.

### Mouse with humanized liver

All animal experiments were performed in accordance and with approval from the NHLBI Animal Care and Use Committee or the Animal Care Committee of the Central Institute for Experimental Animals (CIEA, Japan). TK-NOG mice, in which a herpes simplex virus type 1 thymidine kinase (TK) transgene under a mouse albumin promoter is expressed within the liver of highly immune-deficient NOG mice, were obtained from Taconic Biosciences. The TK converts an antiviral medication ganciclovir (GCV) into a toxic product that allows selective elimination of TK positive cells in vivo. The cryopreserved primary human hepatocytes were obtained from Lonza. The humanized TK-NOG mice were prepared as previously described ^25^. Briefly, The TK-NOG mice at 8-9 weeks old received an i.p. injection of GCV at a dose of 25 mg/kg. One week later, 50ul volume of 1×10^6^ human primary hepatocytes suspended in HBSS solution were transplanted via intra-splenic injection. 8-12 weeks after transplantation, the serum human albumin in the mice were measured as an index of the extent of human hepatocytes replacement. Humanized TK-NOG mice with serum human albumin levels above 0.5 mg/ml were used for experiments, in which human hepatic genes could be reliably detected by q-PCR. For the fasting-refeeding study, humanized mice were produced and the experiment was carried out at CIEA. Humanized mice for the rest of the study were produced and analyzed at NHLBI. For the fasting-refeeding study, humanized TK-NOG mice were allowed free access to food (Fed) or subjected to a twenty-four hours food withdrawal (Fasting) or subjected to a twenty-four hours food withdrawal followed by a four-hours refeeding (Refeeding) before tissue harvest. Animal data were excluded from experiments based on pre-established criteria of visible abnormal liver structure during sample harvest or other health issues such as fighting wounds or infections. According to the variability of metabolic parameters, group size was determined based on previous studies using similar assays within the laboratory and pilot experiments. Experimenters were not blinded to treatment group.

### RNA extraction, RNA-seq and quantitative real-time PCR analysis

Total RNA was isolated from liver tissues using Trizol reagent (Invitrogen). After Turbo DNA-free DNase treatment (Ambion), the construction of strand specific sequencing libraries using illumina TruSeq RNA sample Prep kit and the sequencing were performed at NHLBI DNA Sequencing and Genomics Core. And the reverse transcription was carried out with SuperScript® III First-Strand Synthesis system (Invitrogen) using 1 µg of RNA. Quantitative real-time RT-PCR was performed on a ViiA™ 7 Real-Time PCR System (Applied Biosystems Inc.) The PCR program was: 2 min 30 s at 95°C for enzyme activation, 40 cycles of 15 s at 95°C, and 1 min at 60°C. Melting curve analysis was performed to confirm the real-time PCR products. For detecting the expressions of human genes in humanized liver samples, human-specific primers were designed and quantitation was normalized to human 16S rRNA levels. For detecting the expressions of mouse genes in regular mice (C57BL/6), 18S rRNA was used as the internal control. For detecting the expression of hLMR1 in human tissues, the Human MTC Panel I (Cat. No. 636742, Takara Bio Inc.) was used. The full primer sequences used are provided as below:

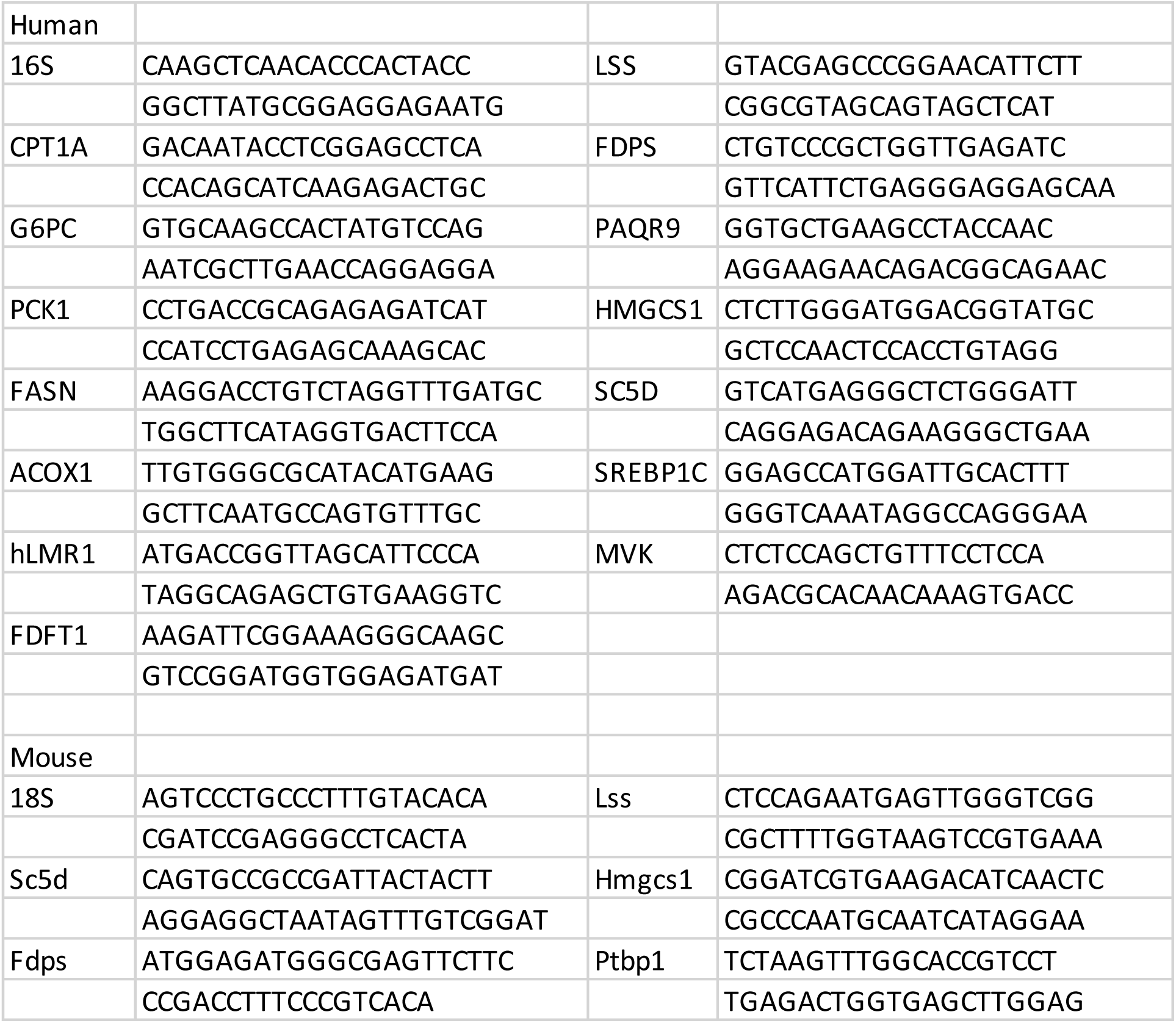

### In vitro translation

The in vitro translation analysis was performed using the TnT® Quick Coupled Transcription/Translation System from Promega by following the manufacture’s protocol. The translated protein was visualized by using IRDye Streptavidin (LI-COR) which detected the biotinylated lysine incorporated into the translated proteins. The open reading frame of yellow fluorescent protein (YFP) was used as a positive control.

### Adenovirus production and in vivo adenovirus administration

The shRNAs for hLMR1 and mouse Ptbp1 were designed using the following sequences (hLMR1 shRNA: CCTTCACAGCTCTGCCTAA; mouse Ptbp1 shRNA: GCACCGTCCTGAAGATCAT). The hairpin template oligonucleotides were synthesized by Integrated DNA Technologies and were subsequently cloned into the adenovirus vector of the pAD/Block-it system (Invitrogen) according to the manufacturer’s protocols. Overexpression construct of hLMR1 was generated by PCR-amplifying the sequence of ENST00000476385.1 using human liver cDNA sample. The sequence was subsequently cloned into pAdv5 adenovirus vector for virus packaging. Adenoviruses were amplified in HEK293A cells and purified by CsCl gradient centrifugation. Purified viruses were desalted with PD10 columns (GE Healthcare Life Sciences) and titered with Adeno-X Rapid Titer Kit (Clontech). Adenoviruses were delivered into mice intravenously at 5 × 10^8^ pfu/mouse for overexpression experiments, 1 × 10^9^ pfu/mouse for knockdown experiments. After seven days, tissue samples were harvested for further analysis.

### CRISPR activation in humanized mice

CRISPR activation assay was conducted using the SAM system (http://sam.genome-engineering.org/protocols/). The sgRNA for hLMR1 was designed using the following sequence (Forward: CACCGGACAGACAGGAGAGCAGACT, Reverse: AAACAGTCTGCTCTCCTGTCTGTCC). Briefly, the template was ligated into the sgRNA (MS2) cloning backbone (Addgene: 61424) using Golden-Gate reaction (SAM) after Golden-Gate annealing. The expression cassettes of dCas9-VP64 (Addgene: 61422) and MS2-P65-HSF1 (Addgene: 61423) were sub-cloned into pAdv5 vector (Invitrogen) for virus packaging. SgRNA elements with the U6 promoter were amplified and subsequently cloned into pAd/PL adenovirus vector (Invitrogen) for virus packaging. Viruses were amplified, desalted and tittered as described above. Three adenoviruses (1:1:1) were delivered into humanized mice intravenously at a total of 5 × 10^8^ pfu/mouse. After seven days, tissue samples were harvested for further analysis.

### RNA pull-down assay

RNA pull-down was performed as described previously^28^. Briefly, biotin-labeled RNAs were transcribed in vitro using the Biotin RNA Labeling Mix and T7 RNA polymerase (Ambion) and purified with the RNeasy Mini Kit (QIAGEN). Folded RNAs (3 ug) were added into 5 mg pre-cleared liver lysates of humanized mice (supplemented with 0.2 mg/ml heparin, 0.2 mg/ml yeast tRNA, and 1 mM DTT) and incubated at 4°C for 1 hr. There were 60 μl of washed Streptavidin-coupled Dynabeads (Invitrogen) that were added to each binding reaction and further incubated at 4°C for 1 hr. Beads were washed five times with lysis buffer (150 mM NaCl, 20 mM Tris pH 7.4, 1 mM EDTA, 0.5% Triton X-100 with Protease/Phosphatase Inhibitor Cocktail and RNaseOUT) and heated at 70°C for 10 min in 1× lithium dodecyl sulfate (LDS) loading buffer, and retrieved proteins were visualized by SDS-PAGE and silver staining. The unique protein bands shown in the hLMR1 RNA pull-down were identified by mass spectrometry analysis at NHLBI Proteomics Facility.

### RNA immunoprecipitation (RIP)

To prepare liver tissue lysates, frozen liver tissues were homogenized using a dounce homogenizer with 15-20 strokes in RIP buffer (150 mM NaCl, 20 mM Tris pH 7.4, 1 mM EDTA, 0.5% Triton X-100 with Protease/Phosphatase Inhibitor Cocktail and RNaseOUT). For each RIP, 5 μg rabbit IgG or PTBP1 Antibody (Cat. No. 32-4800, FISHER SCIENTIFIC) were first incubated with 30 μl washed Dynabeads® Protein G in 300 μl RIP buffer supplemented with 0.2 mg/ml BSA, 0.2 mg/ml Heparin and 0.2mg/ml *EcoRI* tRNA for one hour. Then the antibody coupled beads were added to 5 mg of liver tissue lysates diluted in 500 μL RIP buffer and incubated for 3 hours at 4°C with gentle rotation. Beads were washed briefly five times with RIP buffer. At the final wash, one fifth of beads were used for protein analysis and the rest of beads were resuspended in 1 ml of Trizol for RNA extraction. Co-precipitated RNAs were isolated and analyzed by RT-PCR.

### Chromatin immunoprecipitation (ChIP) analysis

ChIP assays of frozen liver tissues of humanized mice were performed using a Simple ChIP Enzymatic Chromatin IP kit (Cell Signaling Technology) according to the manufacturer’s protocol. Immunoprecipitation was performed using RNA Poly II ChIP validated antibody (MilliporeSigma, Cat: 17-620), PTBP1 Antibody (FISHER SCIENTIFIC, Cat. No. 32-4800) or with rabbit IgG as a negative control. For RNA poly II ChIP, the DNA in each ChIP were determined by quantitative real-time PCR analysis using primers amplifying the genomic sequences covering the transcriptional start sites of genes. For PTBP1 ChIP, the DNA in each ChIP were determined by quantitative real-time PCR analysis using primers amplifying the genomic sequences covering the promoters of genes. The primers used are list as below. The relative enrichment was calculated by normalized the amount of ChIP DNA to input DNA and compared with IgG control as fold enrichment.

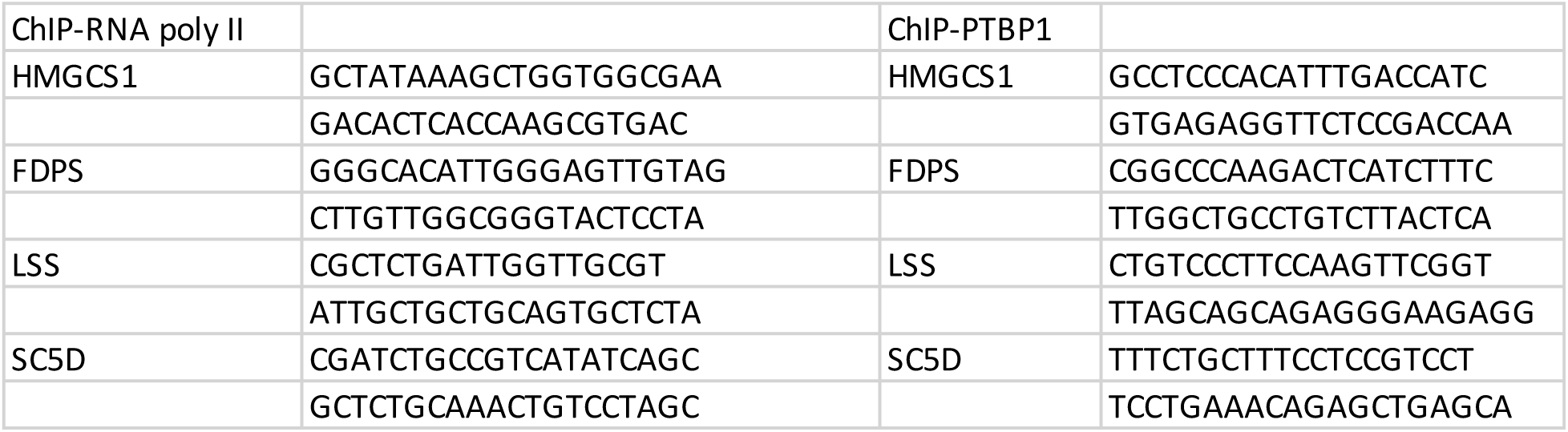

### Luciferase Reporter Assay

The human HMGCS1 promoter was amplified (forward primer: GTCCATCGGAATTAGTTTAGCCTGTGC, reverse primer: CAATCGCGGCCGGTAGAGTTG) and cloned into the pGL3-Basic Vector (Promega). Full-length PTBP1 expression vector and control vector were purchased from OriGene (Cat: RC201779 and PS100001). The HEK293A cells were maintained in DMEM supplemented with 10% Cosmic Calf Serum. Cells were transfected with pGL3-HMGCS1 promoter, the PTBP1, pAd-hLMR1 or control vectors using Lipofectamine 2000 (Invitrogen), and luciferase assays were performed 24 hr later using the Dual-Luciferase Reporter Assay Kit (Promega). Transfection efficiency was measured by normalization to Renilla luciferase activity expressed from a co-transfected pTK-RL vector (Promega).

### Immunoblotting

For Immunoblotting analyses, the cells and tissues were lysed in 1% SDS lysis buffer containing phosphatase inhibitors (Sigma) and a protease inhibitor cocktail (Roche). The lysate was subjected to SDS–PAGE, transferred to polyvinylidene fluoride (PVDF) membranes, and incubated with the primary antibody followed by the fluorescence conjugated secondary antibody (LI-COR). The bound antibody was visualized using a quantitative fluorescence imaging system (LI-COR). The PTBP1 Antibody (Cat. No. 32-4800) was from FISHER SCIENTIFIC.

### Measurement of lipid levels in liver tissues and plasma

The liver and plasma cholesterol levels in regular mice were measured by using Cholesterol Assay Kit from Abcam (ab65390) and normalized to tissue weights. To measure human LDL-VLDL cholesterol levels in the plasma of humanized mice, human Apolipoprotein B containing lipoproteins in the plasma of humanized mice were immunoprecipitated by using LipoSep IP™ reagent (Sun Diagnostics, Cat: LS-01). The immunoprecipitated pallets were first washed with PBS, and then were resuspended in PBS plus 0.5% NP-40 to release lipids from lipoproteins. After a brief centrifuge, the cholesterol levels in supernatant were measured by using Cholesterol Assay Kit from Abcam (ab65390) (as in Supplemental Figure 1D), or further normalized to the humanized ratio of each mouse (as in Figure 4D). The humanized ratio was determined by the relative expression levels of human 16S in the real-time PCR analyses using cDNA prepared from the homogeneous powder of each humanized liver tissue.

---

**Figure S1 related to Figures 3 and 4.**
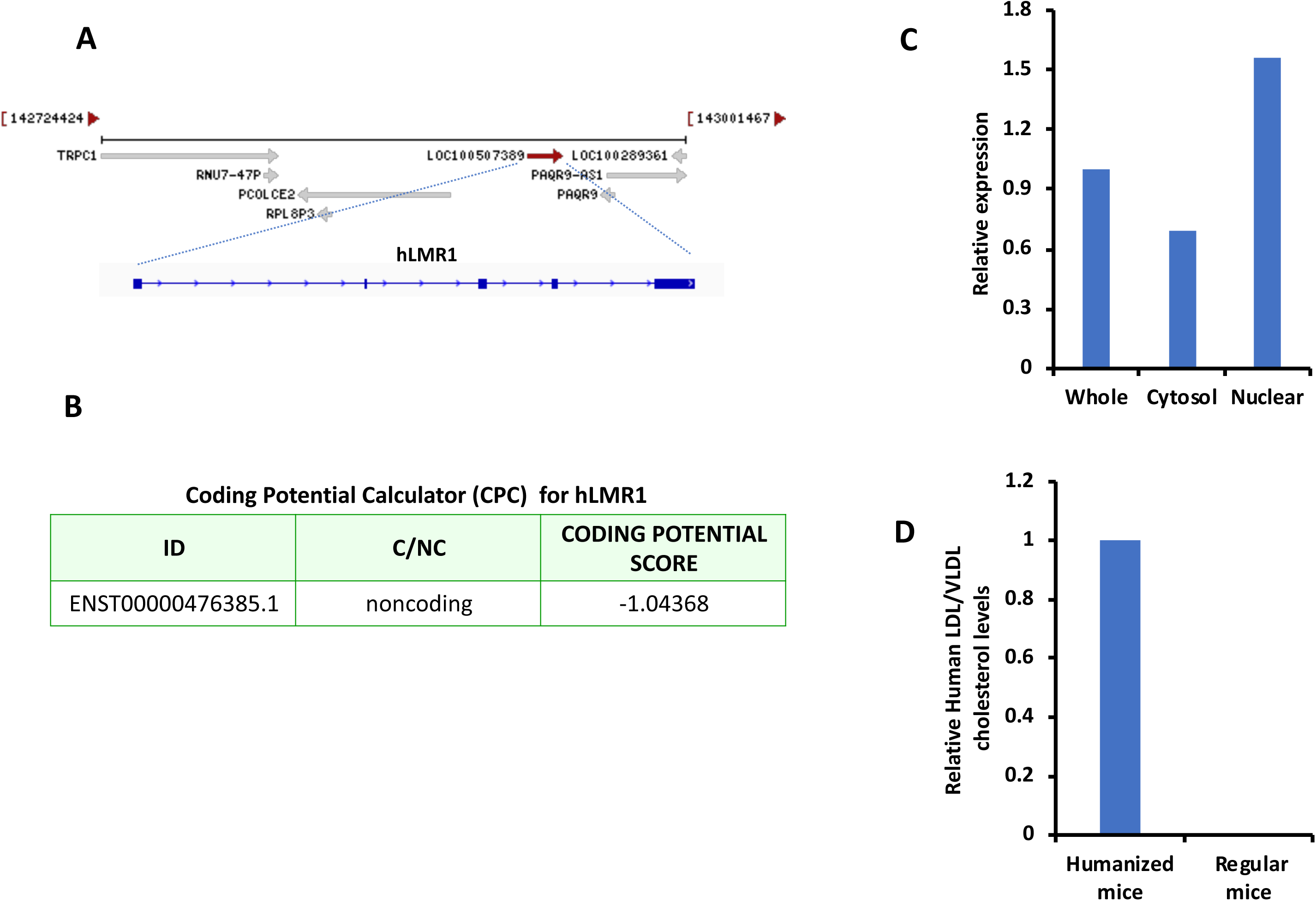
(A) Gene structure of hLMR1 in human genome. (B) Coding potential analysis for hLMR1 using Coding Potential Calculator (CPC). (C) Cellular localization analysis of hLMR1. (D) Relative cholesterol levels in human ApoB-containing lipoproteins purified from the plasma of humanized mice or regular mice.

**Table S1 Related to Figure 2**

List of the genes in the top quartile of gene expression variabilities.

**Table S2 Related to Figure 2**

List of the differentially expressed genes in human NAFLD studies.

**Table S3 Related to Figure 2**

List of the differentially expressed genes in human LCD intervention studies.

**Table S4 Related to Figure 2**

List of the hLMRs and their associated GO terms.

**Table S5 Related to Figure 3**

List of the differentially expressed genes in fasting-refeeding studies of humanized mice.

